# Power and precision: Evaluation and recommendations of quantitative trait analysis methods for RNA expression levels in the Hybrid Rat Diversity Panel

**DOI:** 10.1101/2022.07.14.500105

**Authors:** Jack Pattee, Lauren A. Vanderlinden, Spencer Mahaffey, Paula Hoffman, Boris Tabakoff, Laura M Saba

## Abstract

The Hybrid Rat Diversity Panel (HRDP) is a stable and well-characterized set of more than 90 inbred rat strains that can be leveraged for systems genetics approaches to understanding the genetic and genomic variation associated with complex disease. The HRDP exhibits substantial between-strain diversity while retaining substantial within-strain isogenicity, allowing for the precise mapping of genetic variation associated with complex phenotypes and providing statistical power to identify associated variants. In order to robustly identify associated genetic variants, it is important to account for the population structure induced by inbreeding. To this end, we investigate the performance of various plausible approaches towards modeling quantitative traits in the HRDP and quantify their operating characteristics. Towards facilitating study planning and design, we conduct extensive simulations to investigate the power of genetic association analyses in the HRDP, and characterize the impressive attained power.

## 1 Introduction

The Hybrid Rat Diversity Panel (HRDP) is large panel of 99 inbred rat strains consisting of three subpopulations: the HXB/BXH recombinant inbred (RI) panel (30 strains), the FXLE/LEXF RI panel (34 strains), and a group of divergent classic inbred strains (35 strains) (Tabakoff et al. 2019). The HRDP was designed to maximize both power and mapping resolution when examining genetic determinants of complex traits, i.e., quantitative trait locus (QTL) analyses. High resolution is facilitated by the inclusion of 35 classic inbred strains, while power is provided by the two recombinant inbred panels. Hybrid diversity panels are a popular framework in model organism analysis; the Hybrid Mouse Diversity Panel (HMDP) has been leveraged in systems genetics approaches to studying various genetic traits including heart failure, plasma lipid levels, and insulin resistance (Lusis et al. 2016). However, the usefulness of panels of inbred mice is somewhat hampered by the similar genetic origin of laboratory mouse strains (Reuveni, Birney, and Gross 2010) and the difficulty of breeding “wild-type” mouse strains with founder strains to derive recombinant inbred strains (Odet et al. 2015). Additionally, rats are often preferred to mice for behavioral and physiologic studies due to their greater size and more distinguishable central nervous system anatomy (Parker et al. 2014). These factors motivate the development and the usage of the HRDP for systems genetics applications.

Despite the large number of SNPs statistically associated with complex traits, the mechanism by which genetic variation affects complex traits is often not well understood. One mechanism for the genetic control of complex traits is the modulation of gene expression by SNPs (Zhang et al. 2015). A substantial proportion of variants associated with complex traits are also associated with variation in gene expression (Brænne et al. 2015). This illustrates the importance of identifying genetic variants associated with variation in transcript levels, as further elucidating the interaction of genetic variation, gene expression, and complex trait presentation is of substantial importance for parsing the genetic etiology of disease. Gene expression can be treated as a quantitative trait in a genome-wide association scan; variants associated with gene expression in such an analysis are termed expression quantitative trait loci, or eQTLs. Whereas behavioral and physiological phenotypes are often influenced by hundreds of loci throughout the genome, variants associated with transcript levels are often located in the coding sequence of the gene or in nearby enhancer or promoter regions. Variants that influence transcript levels that are in or near the coding region of the gene are often called cis-eQTLs, although a true cis mechanism of control would need to be validated with additional studies. More distant variants are termed trans-eQTLs. It is thought that the identification of cis-eQTLs can illuminate a potential mediation mechanism for the effect of genetic variants on behavioral and physiological phenotypes, whereby variants influence gene expression which in turn affects behavioral and physiological traits (Gusev et al. 2016).

Analyzing QTLs in a panel of inbred organisms with complex relationships among individuals (such as the HRDP) requires modeling considerations for the effect of population structure. Initial analyses of the HMDP were substantially confounded by failure to properly account for the effect of population structure, resulting in poor type I error control (Kang et al. 2008). Linear mixed models with random effect structure dictated by the estimated genetic relationship matrix have demonstrated good performance in the analysis of eQTLs in populations with shared ancestry. A popular implementation of linear mixed models of this type for the analysis of genetic data is implemented in the software GEMMA (Zhou and Stephens 2012).

This paper examines four methodological approaches towards modeling eQTLs in the HRDP. In particular, we analyze a 43-strain subset of the HRDP for which we have both RNA expression levels and genotype information. This 43-strain subset consists of 30 strains from the HXB/BXH RI panel as well as 13 classic inbred strains. The four methods we compare are ordinary least squares linear regression, linear mixed modeling, linear mixed modeling with leave-one-chromosome-out (LOCO) kinship matrix calculation, and linear mixed modeling by subpopulation followed by meta-analysis. All linear mixed models described in this study were estimated via GEMMA. We compare the performance of these four methods in real data application and in simulation with the goal of arriving at a modeling recommendation for eQTL analyses of the HRDP data. Our analysis indicates that linear mixed modeling via GEMMA is appropriate for eQTL analysis of the HRDP data. Implementing LOCO for kinship calculation increase power but may also slightly inflate type I error (i.e., false positives). We additionally conduct a simulation study with 92 HRDP strains with genotype information and empirically relate QTL heritability to power for the three component subpopulations and the full HRDP. This empirical power analysis may inform study design and sample size calculations for researchers interested in using the HRDP.

## 2 Materials and Methods

### 2.1 Animals

The HRDP is large panel of 99 inbred rat strains consisting of three subpopulations: the HXB/BXH recombinant inbred (RI) panel, the FXLE/LEXF RI panel and a group of divergent classic inbred strains (Tabakoff et al. 2019). We obtained brain tissue from a subset of the HRDP (45 strains). Rats from HXB/BXH RI panel (Pravenec et al. 1989) including its progenitor strains (BN-Lx/Cub and SHR/OlaIpcv) used for RNA-Seq analyses were maintained by Dr. Michal Pravenec at the Institute of Physiology of the Czech Academy of Sciences. For this RI panel, the University of Colorado Anschutz Medical Campus received shipments of brain tissue from male rats (~70-90 days old) stored in liquid nitrogen. The process of retrieving tissues was performed in accordance with the Animal Protection Law of the Czech Republic and approved by the Ethics Committee of the Institute of Physiology, Czech Academy of Sciences, Prague. The other 13 inbred strains from the group of divergent classic inbred strains were received at the University of Colorado Anschutz Medical Campus from either Charles River or Envigo. Like the HXB/BXH RI panel, male rats (~70-90 days old) were sacrificed via CO2 exposure and quickly decapitated. Brains were cut in half sagittally, placed immediately in RNAlater, and stored in a −80° C freezer.

### 2.2 DNA Sequence Variants

A genetic marker set (specifically SNPs) from the STAR consortium (http://www.snp-star.eu/; STAR Consortium et al 2008) was used for QTL analyses. Probe sequences from the original arrays were aligned to the RN6 version of the rat genome using BLAT (Kent 2002) (v2.7.6). SNPs were retained if their probe sequence aligned both perfectly and uniquely to the rat genome. Of the 99 strains in the HRDP, 92 strains had genotype information (43 of the 45 with brain RNA expression). Strains were matched by substrain (when an established inbred strain genetically diverges into two separate populations for reasons such as location of maintenance and breeding) when possible. SNPs were further eliminated if more than 10% of the 92 HRDP were missing a genotype call. To identify potential genotype errors or SNPs misplaced in the genome, genomic maps were estimated separately for the HXB/BXH recombinant inbred strains and for the FXLE/LEXF recombinant inbred strains using the qtl package in R (Broman et al. 2003; version 1.50). SNPs with “improbable recombination rates” were identified by a distance of 20 cM or more between the SNP and both of its adjacent SNPs in the estimated genomic maps. If a SNP was the first or last SNP on a chromosome and had 20 cM between it and the closest SNP on the same chromosome, it was also eliminated. SNPs identified as having an improbable recombination rate in either RI population were eliminated.

### 2.3 Brain RNA Expression Levels

Total RNA was isolated from whole brain tissue from 3 biological replicates per strain for 45 strains of the HRDP using QIAzol (Qiagen, Valencia, CA, USA). The brain was split sagitally and half of the brain was used for RNA sequencing. The RNAeasy Plus Universal Midi Kit (Qiagen) was used to separate long (>200 nucleotides) and short (<200 nucleotides) fractions. The long RNA fraction was purified using the RNeasy Mini Kit (Qiagen). Sequencing libraries for the long RNA fraction were constructed using the Illumina TruSeq Stranded Total RNA Sample Preparation Kit with Ribo-Zero ribosomal RNA reduction chemistry (Illumina) in accordance with the manufacturer’s instructions with the exception that for later batches multiple Ribo-Zero washes were used to reduce the amount of rRNA sequenced. Samples were sequenced in eight batches on an Illumina HiSeq2500 or HiSeq4000 (Illumina, San Diego, CA, USA) in High Output mode to generate 2×100 or 2×150 paired end reads.

Prior to alignment, reads were demultiplexed and read fragments were trimmed for adaptors and for quality using Cutadapt (Kechin et al. 2017) (version 1.9.1). Reads were eliminated if the trimmed length of either read fragment was less than 20 nucleotides. Reads aligned to ribosomal RNA from the RepeatMasker database (Smit, Hubley, and Green 1996) (accessed through the UCSC Genome Browser; https://genome.ucsc.edu/) were also eliminated. This alignment was done using Bowtie 2 (v.2.3.4.3) (Langmead and Salzberg 2012). The remaining reads were aligned to the RN6 version of their respective strain-specific genomes containing only SNPs derived from our DNA sequencing (Saba et al. 2015) using HISAT2 (Kim, Langmead, and Salzberg 2015) (v.2.1.0) with the default settings and samtools(v1.3) mpileup variant calls. A genome- and transcriptome-guided reconstruction was executed for each strain separately using the StringTie (Pertea et al. 2015) algorithm and software (version 1.3.5). The Ensembl Rat Transcriptome (version 96) was used to guide the reconstruction process. The strain-specific reconstructed transcriptomes were merged using the StringTie merge function. The RNA-Seq by Expectation Maximization (RSEM) algorithm (Li and Dewey 2011) (v1.2.31) was used to estimate the read coverage of each isoform and each gene within the reconstructed transcriptome. Within the reconstructed transcriptome, high quality isoforms were identified using two criteria: 1) detection above background (i.e., an estimated read count of at least 1 in at least 2/3 of the samples) and 2) an effective length greater than or equal to 200 nucleotides. Isoforms that did not meet these criteria were eliminated and the entire transcriptome was re-quantitated for each sample to allow for the redistribution of reads that aligned to multiple isoforms in the context of the reduced transcriptome.

After the second quantitation, the final count matrix used for eQTL analyses was generated by: 1) eliminating the few isoforms that were not detected above background, as defined above, after re-quantitation, 2) eliminating individual samples with low sequencing efficiency (i.e., < 10 million estimated read counts from RSEM), 3) eliminating individual samples when a few transcripts dominate the read count (i.e., one fourth or more of a sample’s isoforms had an estimated read count of zero), and 4) eliminating extreme outliers. Extreme outliers were systematically identified by examining all pairwise correlations among a sample and all other samples using read counts that had been transformed using a regularized log (rlog) (Love, Huber, and Anders 2014). If more than 50 of the correlation coefficients estimated for a single sample were less than 0.90, that sample was removed.

The final count matrix was adjusted for batch effects and other technical effects using the first factor from an RUV (Removing Unwanted Variance) (Risso et al. 2014) analysis based empirical negative control isoforms. Negative control isoforms were identified in a differential expression analysis using DESeq2 (Love, Huber, and Anders 2014) to test for association between isoform expression and strain. Negative control isoforms were within the least significant quartile of the data and also had an average counts per million value greater than 0. The ‘adjusted’ read counts were transformed using the regularized log function (Love, Huber, and Anders 2014). This final transformed data was used in all subsequent statistical analyses.

### 2.4 QTL Analyses

This paper compares four methods for the analysis of QTLs. These models are as follows: 1) a linear regression model, 2) a linear mixed model, 3) a linear mixed model with leave one chromosome out kinship matrix calculation, and 4) a linear mixed model stratified by subpopulation followed by meta-analysis.

Linear regression, where each SNP is pairwise associated with the expression levels of each isoform, is often used for eQTL analyses (Michaelson, Loguercio, and Beyer 2009) (Veyrieras et al. 2008) when samples are equally related to one another (i.e., are independent and identically distributed). This model entails ordinary least squares regression with a single SNP as the independent variable and a single RNA expression phenotype as the dependent variable. The ordinary least squares model makes several strong assumptions about the structure of the residual error, some of which may be violated in our context. For example, it may be that the relatedness between strains in the HRDP data violates the sphericity assumption of the model, which assumes that the error terms are uncorrelated with one another. If this assumption is violated, the linear regression model may experience type I error inflation (Kang et al. 2008) (Cervino et al. 2007). Linear regression analyses were conducted in R version 3.6.1 using the R package ‘qtl’ version 1.46-2 (Broman et al. 2003).

Linear mixed models are a popular tool for conducting eQTL analyses in the presence of confounding by shared ancestry (Zhou and Stephens 2012) (Kang et al. 2008). The linear mixed model allows for the presence of correlation between the RNA expression phenotypes of related strains by introducing a random effect into the model. The distribution of this random effect is assumed to be normal, with error variance proportional to the kinship matrix. The kinship matrix, which can be estimated from genetic data, approximates the degree of genetic relatedness between strains in the study. Our implementation of the linear mixed model is via Genome-wide Efficient Mixed Model Association (GEMMA) (Zhou and Stephens 2012). All GEMMA models were fit using GEMMA version 0.98.1 for Linux.

An adjustment to the linear mixed model as described above is the linear mixed model with leave-one-chromosome-out (LOCO) kinship matrix calculation, henceforth GEMMA LOCO. This approach has some theoretical benefits as compared to the standard GEMMA approach (Yang et al. 2014). An assumption of the linear mixed model is that fixed effects, which in our context is the SNP effect of interest, are uncorrelated with the random effect. By excluding SNPs that are on the same chromosome as the fixed effect SNP from the calculation of the kinship matrix, we ensure that there is no correlation between the fixed and random effects in the model. This approach increases the likelihood that the assumptions of the linear mixed model are met.

Given that the HRDP sample in our study is a heterogeneous sample of two distinct subpopulations, it may be appropriate to conduct a meta-analysis of the two subpopulations. This would entail estimating linear mixed models for the two subpopulations separately and combining results (i.e., p-values) through a meta-analysis approach. For this strategy, we estimate separate GEMMA models for the recombinant inbred population and the classic inbred population and combine the results using Stouffer’s method (Stouffer et al. 1949). The progenitors of the HXB/BXH RI panel were included with the classic inbred population rather than the RI population.

All p-values from linear mixed models were calculated via likelihood ratio statistics.

The kinship matrix was estimated using all 18,342 SNPs in the HRDP data. Kinship coefficients are equal to the proportion of shared alleles. Because the set of 18,342 SNPs contains some SNPs with missing observations, the kinship coefficient between two strains is equal to the proportion of shared alleles among all SNPs with non-missing entries for both strains. SNPs were only included as fixed effects in one of the above four models if they passed quality control, which required SNPs to have minor allele frequency > 0.1 and <5% missingness among the strains included in the eQTL analysis. If a meta-analysis was performed on subpopulations, only SNPs that passed quality control in both subpopulations were included in the analysis.

The four methods were characterized by their type I error rate (i.e., false positives) and by the number of significant eQTL identified. We cannot directly characterize the type I error rate as the true causal SNPs are unknowable in real data. Thus, we instead use the genomic inflation factor to assess the type I error rate. The genomic inflation factor is defined as the ratio of the median observed χ^2^statistic across all SNPs in a GWAS to the median of the 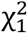 distribution. When interpreting the genomic inflation factor, we assume that only a few SNPs make a substantial contribution to difference in the RNA expression levels of a transcript. We compare the number of cis-SNPs identified as significant by the four methods and use these results in conjunction with the results of the genomic inflation factor analysis to evaluate the performance of the four approaches.

### 2.5 Heritability and Modality

The coefficient of determination (R^2^) from a one-way ANOVA with strain as the predictor and transformed isoform expression as the outcome was used to estimate broad-sense heritability for each isoform. To derive a null distribution of the broad-sense heritability, strain labels were permuted and a distribution of resulting heritabilities were compared to observed distribution. In addition to a broad-sense heritability that does not include information about population structure, SNP-based heritability was estimated to be the proportion of variance explained (PVE) estimated via GEMMA (Zhu and Zhou 2020).

To further understand the genetic architecture of RNA expression traits in the HRDP data, we analyzed the distribution of the RNA expression phenotypes. Motivating this analysis is the idea that highly heritable RNA expression phenotypes may be driven primarily by one or a few strong SNP effects, and these strong SNP effects may induce bi- or multi-modality in the distribution of the RNA expression phenotype, i.e., it may resemble a mixture of two or more normal distributions. Alternatively, RNA expression phenotypes may have many small contributing SNP effects from across the genome, or they may be primarily driven by environmental effects or random variance. In either of these cases, we would expect the distribution of the phenotype to be unimodal.

Modality was assessed by fitting univariate normal mixtures to the RNA expression estimates from 43 strains for a single isoform using the ‘mclust’ R package (Scrucca et al. 2016) version 5.4.7. Three separate models were fit to each isoform that assume that the expression levels are generated from a normal distribution, a mixture of two normal distributions, or a mixture of three normal distributions (i.e., one, two, or three clusters) assuming equal within-cluster variances. The optimal number of clusters (also referred to as ‘modality’) from among one, two and three was selected by choosing the model with the minimum Bayesian information criterion statistic. The relationship between modality and SNP-based heritability was assessed graphically and via ANOVA.

### 2.6 Simulation Study

In conjunction with real data analysis, we conduct a simulation study to assess the type I error and the power of our eQTL analysis approaches. An outline of the simulation setup is detailed here; full details are in Supplementary S1. Simulated data was generated using R version 3.6.3. Broadly, simulations were conducted for distinct populations: those 43 HRDP strains for which we currently have RNA expression phenotypes and genotype information and which comprised the real data analysis, and the full set of 92 HRDP strains for which we have genotype data. These will be referred to as the ‘43 strain simulations’ and the ‘subpopulation simulations’. For the 43 strain simulations, we compare the performance of standard GEMMA, GEMMA LOCO, and the subpopulation meta-analysis. For the subpopulation simulations, we only consider GEMMA LOCO.

The simulation was designed such that the simulated data mimics the real data as closely as possible. To this end, we conduct the simulation as follows. We used real genotype data from the HRDP and simulated continuous RNA expression phenotype data. The phenotype data is generated as a linear combination of SNP effects plus a normally distributed random error term. SNP effects are simulated from a point-normal distribution. The heritability of each simulated RNA expression phenotype is defined as the heritability of the RNA expression phenotype in the real data, where the heritability of the RNA expression phenotype in real data is estimated by the GEMMA PVE.

The modality of the RNA expression phenotype in the real data informs the simulation as well. RNA expression phenotypes estimated to be unimodal in real data were simulated to have a unimodal distribution in the following way. SNP effects from across the genome were simulated from a point-normal distribution. This simulation structure does not prescribe a larger effect size for cis-SNPs than trans-SNPs, thus making it improbable that a single simulated SNP effect would be large enough to induce bimodality. This simulation setup is referred to as ‘no cis effect’. Note that, throughout the paper, we define ‘cis-SNPs’ as those SNPs within 1 Mb of the coding region of the gene. Alternatively, RNA expression phenotypes that were estimated to be bimodal in the real data were simulated to be bi (or multi) modal. This was done by simulating cis-SNP effects from a distribution with larger variance than SNP trans effects. Additionally, the proportion of heritability attributable to cis-SNPs (defined as SNPs within 1Mb of the coding region) for a simulated bimodal RNA expression phenotype was fixed to be between 0.7 and 0.9 of the total SNP-based heritability. This simulation structure was likely to generate one (or few) SNPs with large effect size, which are more likely to induce bimodality in the eQTL phenotype. Simulations were conducted with exactly one nonzero SNP effect per cis region (‘one cis effect’) and a random number of nonzero SNP effects per cis region, with a mean of two nonzero effects (‘multiple cis effects’).

## 3 Results

### 3.1 Genetic marker set

Of the original 20,283 array probes from STAR Consortium SNP dataset (STAR Consortium et al. 2008), 19,391 had a probe sequence that perfectly and uniquely aligned to the RN6 genome with no mismatches. There were 18,342 SNPs in this marker set after eliminating SNPs with more than 10% of genotypes missing among the 92 strains and SNPs with improbable recombination rates. This set of SNPs was used to estimate the kinship matrices for the standard GEMMA and GEMMA LOCO analyses. Only SNPs with minor allele frequency > 0.1 and missingness rate <5% were included as fixed effects in the QTL mapping methods. For the standard GEMMA, GEMMA LOCO, and linear regression approaches, 13,455 SNPs were retained. For the subpopulation meta-analysis approach, 5,797 SNPs were used. There are fewer SNPs in the subpopulation meta-analysis because SNPs were required to pass the MAF and missingness thresholds in both subpopulations, which was a more stringent criterion.

### 3.2 RNA Expression Values

The original reconstruction of brain transcripts produced 568,269 putative transcripts (i.e., isoforms) from 175 RNA-Seq libraries. Sixteen libraries were eliminated due to poor quality. After limiting to only transcripts that were greater than 200 nucleotides in length and transcripts that had an estimated read count of at least 1 in at least 2/3 of the samples, 170,595 transcripts remained and were requantitated using RSEM. After this requantitation, 99,376 genes (all reads aligned to isoforms of this gene are summed) met the criteria of expression above background (estimated read count of at least 1 in at least 2/3 of the samples) and 135 samples met our quality control standards both prior to and after normalization, transformation, and batch correction.

### 3.3 Kinship

Observed kinship values ranged from 0.33 to 0.99 (Figure 1). Of the lowest 5 kinship values, BN-Lx/Cub is one of the strains involved, which resembles the results of the STAR Consortium (STAR Consortium et al. 2008) and the results from full genome sequences of several of the inbred strains (Hermsen et al. 2015). Some very high kinship values were observed, with the highest being 0.9997 between substrains LEW/Crl and LEW/SsNHsd. The pair of strains SHR/OlaIpcv and SHRSP/Crl, which is known to be highly related based on breeding history, has a kinship value of 0.92 (Supplementary Figure 1).

**Figure 1.**
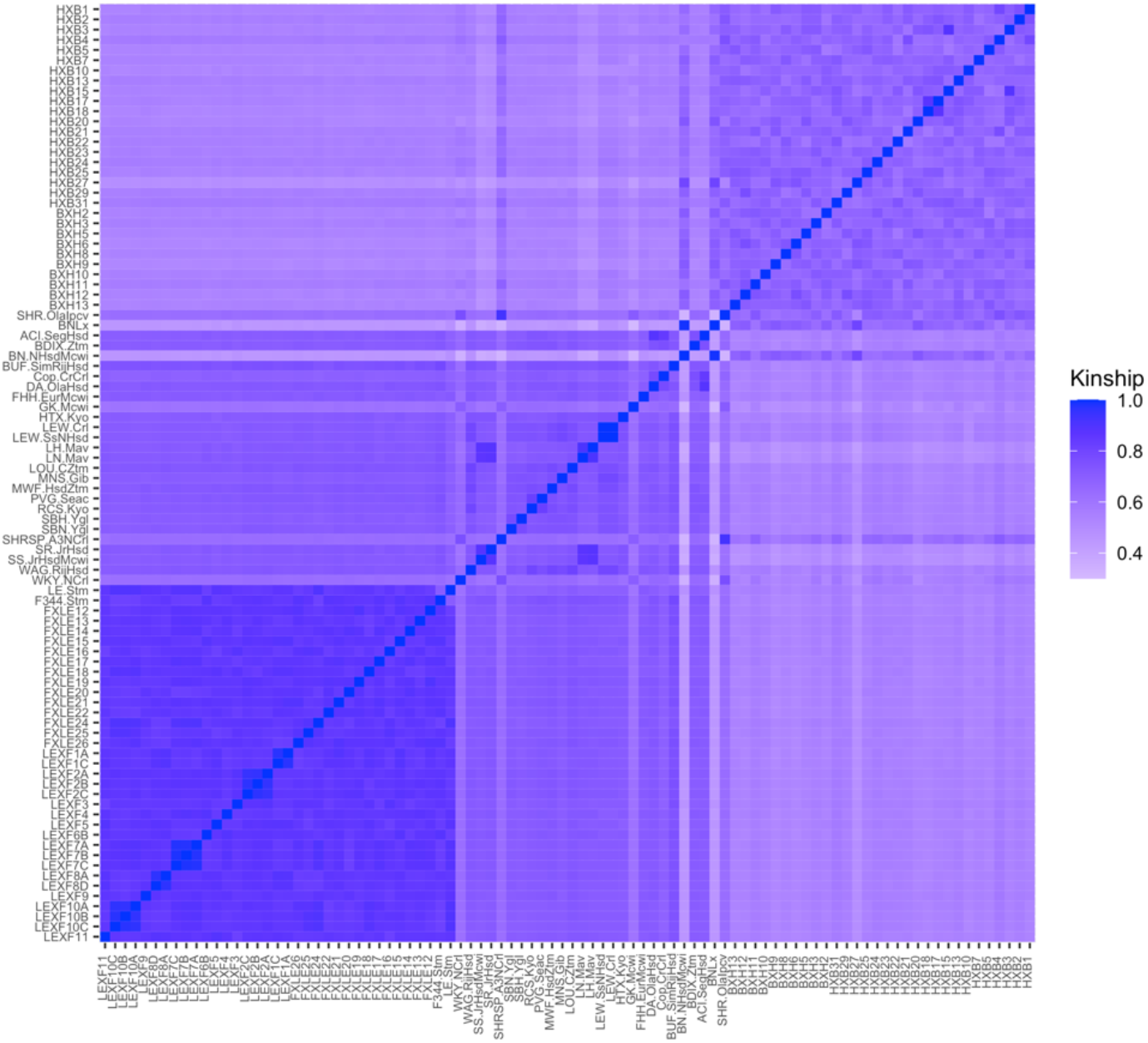
Kinship coefficients for each pairwise genetic relationship between strains in the Hybrid Rat Diversity Panel (HRDP). The upper right block corresponds to the HXB/BXH recombinant inbred panel, while the lower left block corresponds to the FXLE/LEXF recombinant inbred panel and the middle to the classic inbred strains. Brighter colors represent strains that are more closely related as evaluated through SNP information from the STAR Consortium.

### 3.4 Heritability and Modality

We compare the distribution of the observed broad-sense heritability (i.e., 1-way ANOVA R^2^) to the distribution of the permuted R^2^ to investigate the genetic effects on the transcriptome. The center of the distribution of the broad-sense heritabilities from the brain transcriptome is significantly higher than the heritabilities based on permuted values (Figure 2). The true median broad sense heritability is 0.42, while the median of the heritabilities based on permuted values is 0.31. Likewise, many of the heritabilities estimated on the true values are higher than 0.50 (23.6%), whereas very few are greater than 0.50 when the data have been permuted (0.1%).

**Figure 2.**
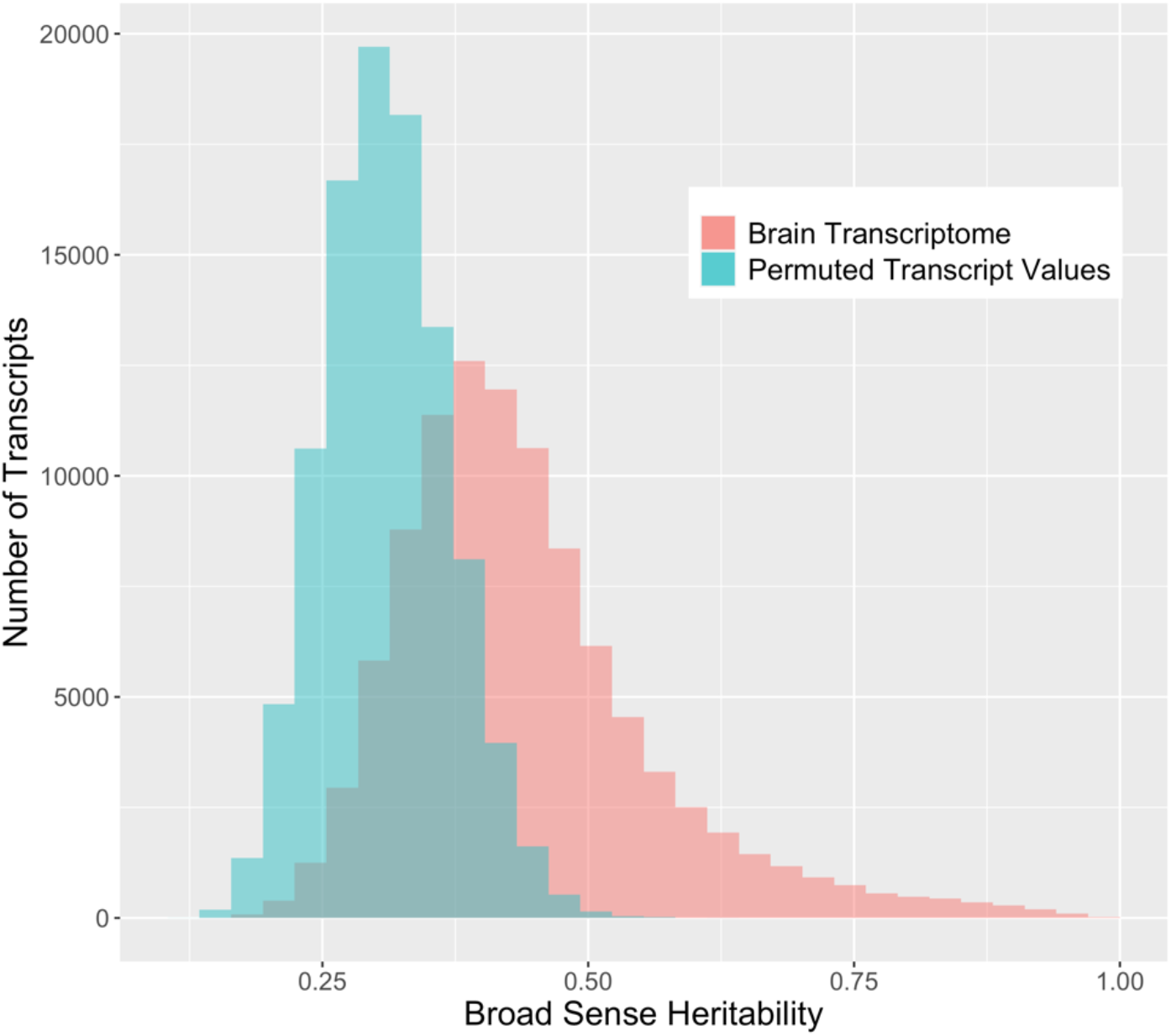
Distribution of broad sense heritabilities for rat brain transcripts compared to permuted expression. The *R*^2^ coefficients estimated via 1-way ANOVA for 99,376 brain transcripts using the true strain labels is compared to permuted expression values. The blue bars represent the distribution of *R*^2^coefficients for the data when permuted to assume no genetic influence, and the distribution of the true brain transcriptome in red represents the broad-sense heritability of RNA expression levels for brain transcripts.

SNP-based heritability for RNA expression phenotypes from brain tissue was assessed via GEMMA (Figure 3). The reader will note a discrepancy between the heritability estimate and displayed in Figure 2 and the heritability estimate displayed in Figure 3, which is due to the following. The GEMMA heritability estimation described in Figure 3 represents the narrow-sense heritability, which is defined as the fraction of phenotypic variance explained by additive SNP effects (Rawlik et al. 2020). Note that narrow-sense heritability estimation is naturally specific to the set of genotyped SNPs. The ANOVA heritability estimation described in Figure 2 is an estimation of the broad-sense heritability, encompasses additive SNP-based heritability and additionally includes the effect of dominance, epistasis, and other heritable trait-determining factors. The majority of transcripts have an estimated narrow-sense heritability for this chip that is relatively small (54.0% with a SNP-based heritability of less than 25%), but a large number of transcripts have a substantial SNP-based heritability.

**Figure 3.**
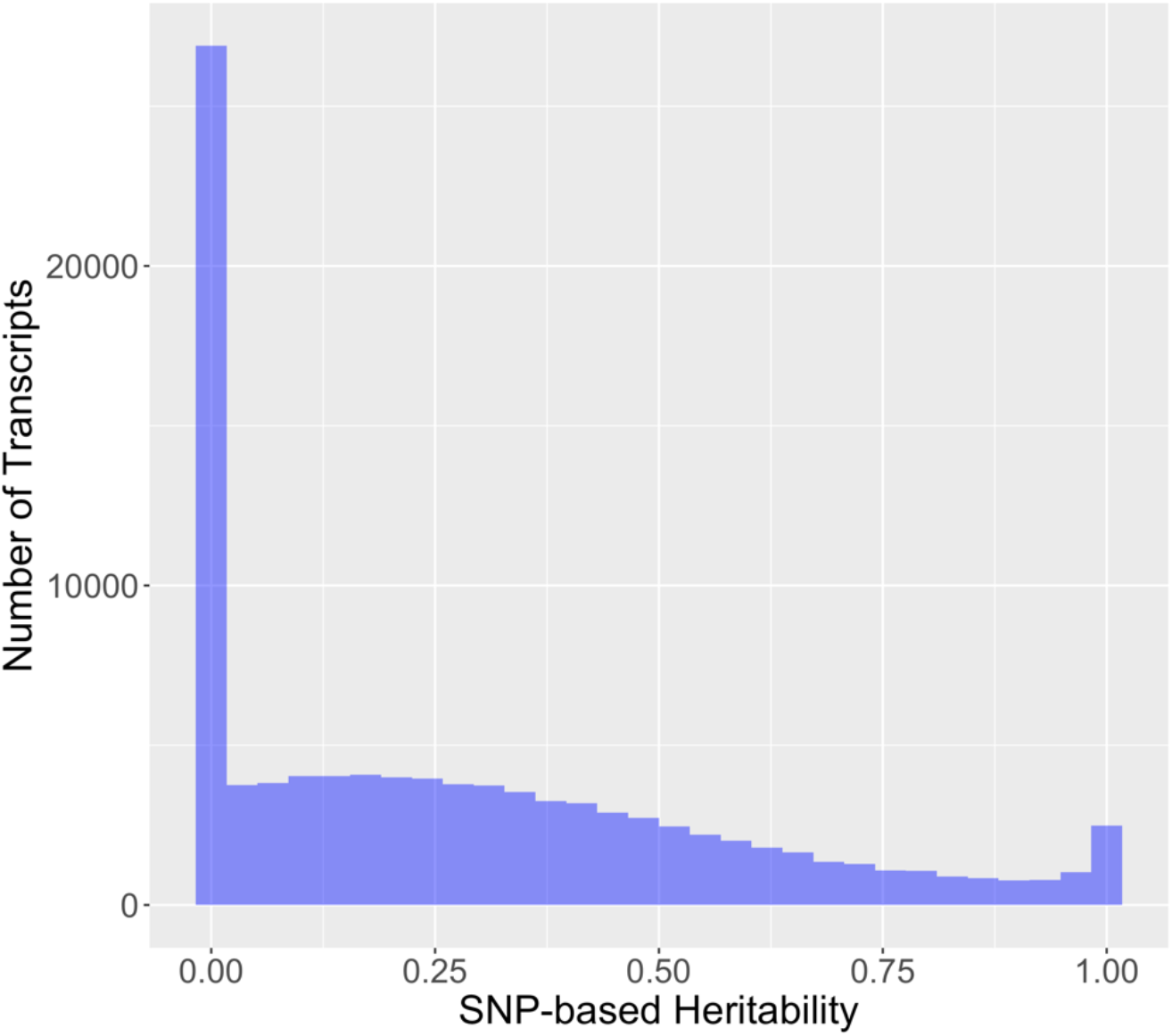
Distribution of SNP-based heritability for 99,376 rat brain RNA transcripts. SNP-based heritability was estimated via the proportion of variance explained from the standard GEMMA modeling using strain means.

Most RNA expression phenotypes (81,992; 82.5%) have a unimodal distribution consistent with many SNPs or non-genetic factors individually contributing a small portion of the variation in expression. A much smaller proportion (15,776; 15.9%) have a bimodal distribution indicative of a single genetic variant with a large effect. A still smaller proportion (1,608; 1.6%) have a trimodal distribution. Modality of the RNA expression levels of a transcript was associated with the SNP-based heritability (omnibus p-value < 1 × 10^−15^, all pairwise comparisons significant). Transcripts with evidence for a bimodal or trimodal distribution of expression values tended to have higher SNP-based heritabilities (Figure 4).

**Figure 4.**
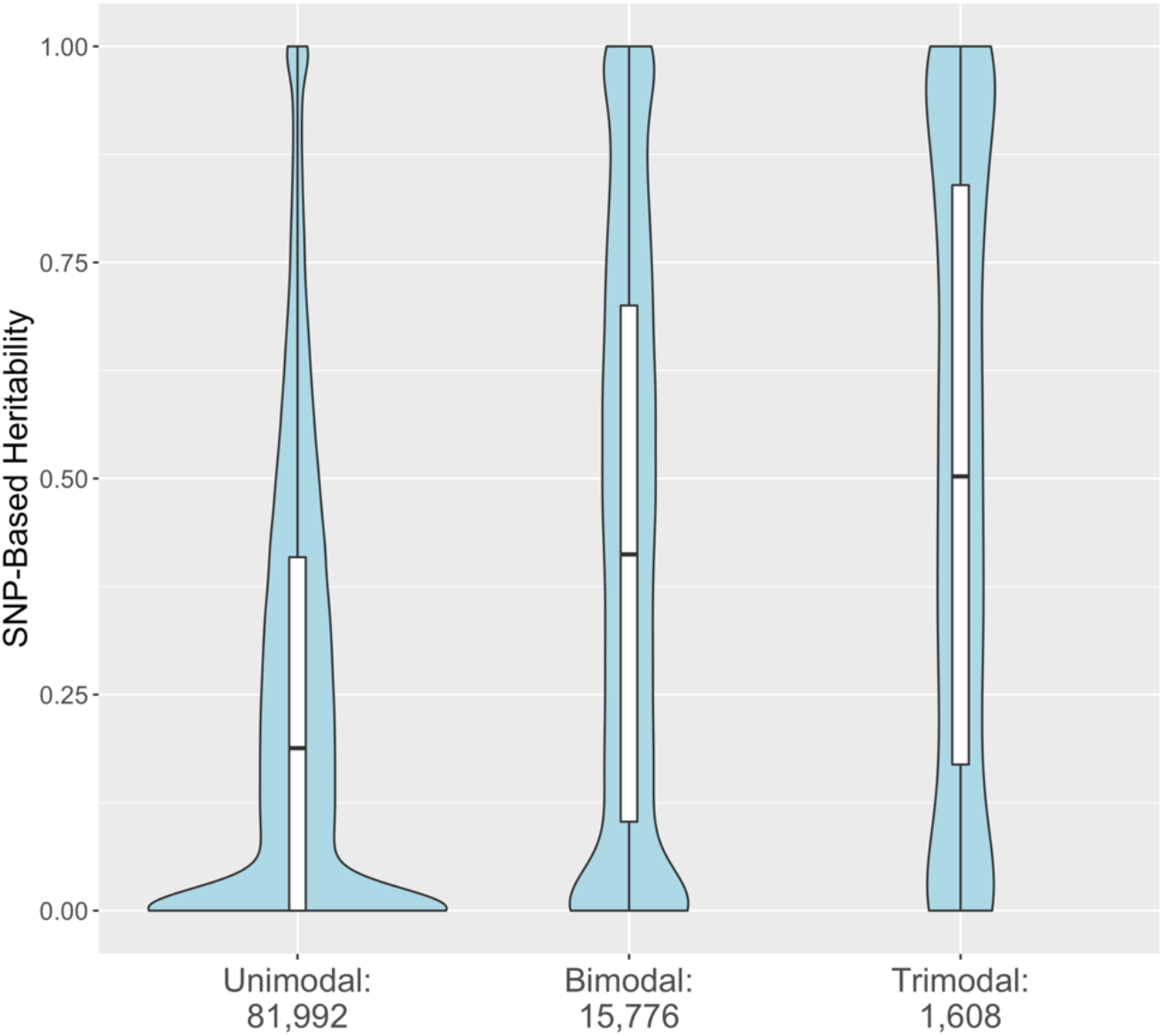
Distribution of SNP-based heritability for 99,376 rat brain RNA transcripts stratified by estimated modality. The SNP-based heritability estimated from strain means and the standard GEMMA model are stratified by the number of modes identified by fitting univariate normal mixtures to the RNA expression estimates. The embedded boxplots display the median and interquartile range.

### 3.5 QTL Results

For each of the four analysis approaches, we calculated genomic inflation factors (GIFs) for each of the 99,376 brain RNA expression phenotypes (Figure 5A). Thus, each method corresponds to a set of 99,376 genomic inflation factors. The linear model generated genomic inflation factors well above the expected value of 1 for all RNA expression phenotypes (Devlin and Roeder 1999); the minimum observed GIF was 1.94. This indicates that every single eQTL phenotype has substantially inflated type I error under the linear regression model. This inflation is likely due to the correlation between RNA expression phenotypes induced by the relatedness among the HRDP strains. The three other methods had similar GIF values (Figure 5B). The subpopulation meta-analysis approach generated a median GIF closest to one (1.09 vs. 1.13 and 1.22 in GEMMA and GEMMA LOCO respectively), but this approach had a bigger spread of values compared to the other two methods (Supplementary Table 1).

**Figure 5.**
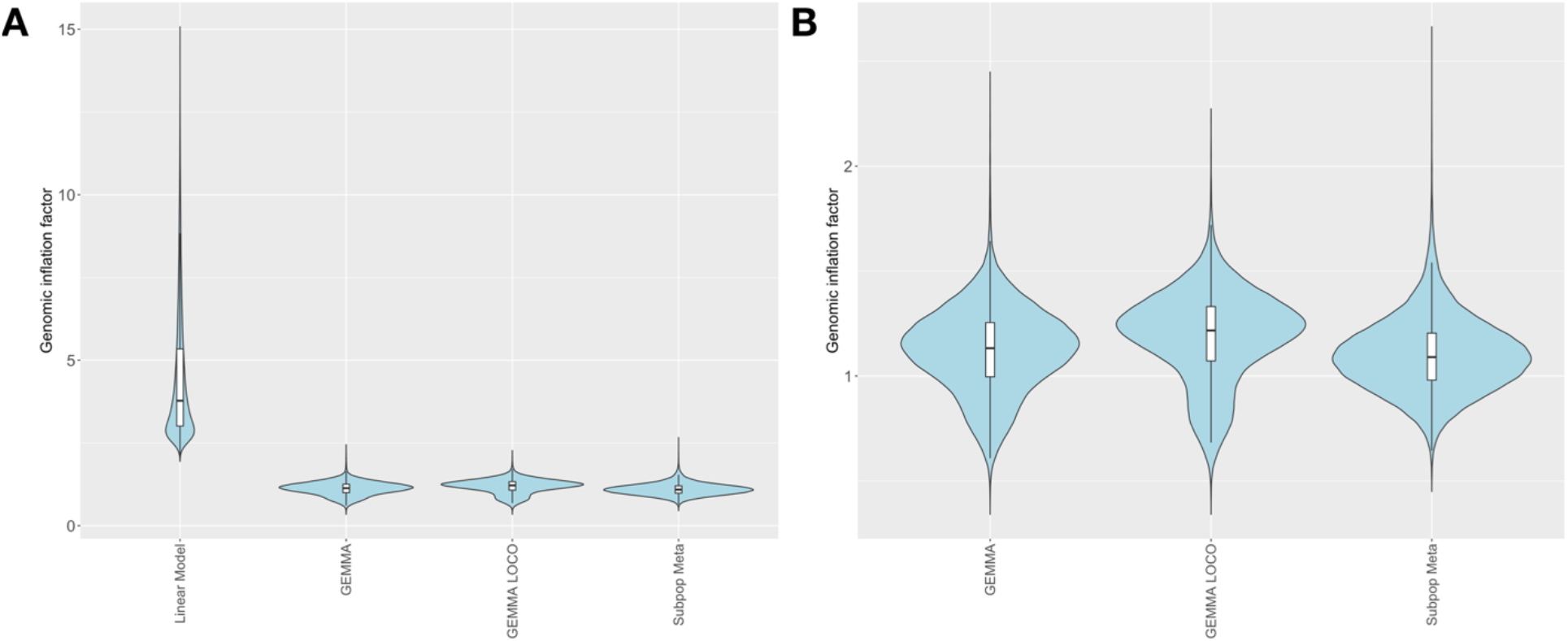
Genomic inflation factors for brain RNA transcripts calculated from p-values derived from different methods of QTL mapping. A) Four mapping methods including linear model, GEMMA, GEMMA LOCO, and subpopulation meta-analysis. B) Three mapping methods with similar ranges (GEMMA LOCO, and subpopulation meta-analysis). The embedded boxplots display the median and the interquartile range.

The number of significant cis-SNPs identified by each of the four approaches was compared. The linear regression model has been invalidated by its inability to control type I error, but we include its results here for completeness. A SNP was determined to be significant if it had a p-value less than 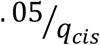, where *q*_*cis*_ is the number of SNPs in the cis-region of the gene (i.e., within 1 Mb of the transcription start or stop site). In this way, we only accounted for the multiplicity among cis-SNPs when determining the multiple testing correction. RNA expression phenotypes without any corresponding cis-SNPs were excluded from the analysis. The number of transcripts with at least 1 significant cis-SNP is similar for the standard GEMMA and GEMMA LOCO approaches, whereas the subpopulation meta-analysis approach identified fewer transcripts with a significant cis-eQTL (Table 1). The dramatic reduction in the number of transcripts with a significant eQTL in the subpopulation meta-analysis approach was mainly due to the reduction in the number of transcripts with a cis-SNP regardless of significance. For the subpopulation meta-analysis approach, a SNP had to have a minor allele frequency greater than 10% in both the recombinant inbred panel and in the panel of classic inbred strains. However, when the proportion of transcripts with a cis-eQTL was calculated based on the number of transcripts with a cis-SNP, the subpopulation meta-analysis still produced a small proportion of cis-eQTL (16.9%) compared to standard GEMMA (18.0%) and GEMMA LOCO (18.9%).

**Table 1.** The number of genes with at least one significant cis-SNP for each of the four methods. The percentages of genes with at least 1 cis-SNP that have a significant cis-eQTL (p-value < 0.05 with a cis-only multiplicity correction) are displayed in column 5. The number of transcripts with at least one cis-SNP (column 4) varies due to more SNPs failing to pass the minor allele frequency cutoff in the subpopulation meta-analysis approach.

| QTL Mapping Method | Transcripts with at least 1 significant cis-eQTL (p-value < 0.05 with a cis-only multiplicity correction) | Transcripts with at least 1 significant cis-eQTL (unadjusted threshold of 10<sup>-5</sup>) | Number of transcripts with at least 1 cis-SNP | Percent of transcripts with at least 1 significant cis-eQTL out of those transcripts with at least on cis-SNP |
| --- | --- | --- | --- | --- |
| Standard GEMMA | 14,618 | 3,627 | 81,184 | 18.0% |
| GEMMA LOCO | 15,186 | 4,035 | 81,184 | 18.7% |
| Subpopulation Meta | 10,886 | 2,506 | 64,542 | 16.9% |
| Linear Model | 24,749 | 6,965 | 81,184 | 30.5% |

The above results, coupled with the reasonable type I error control of the standard GEMMA and GEMMA LOCO methods, is evidence that the subpopulation meta-analysis approach is underpowered. Supplementary Figures 2, 3, and 4 depict the relationship between genes identified by the three approaches. The standard GEMMA and GEMMA LOCO generate fairly similar results, and both agree to a moderate degree with the subpopulation meta-analysis approach.

### 3.6 Simulation Results

The simulation study facilitates the empirical determination of the type I error rate and the power of the methodological approaches in eQTL analysis of the HRDP. Because the linear model was ruled out by the real data analysis, our simulation study only compared the performance of standard GEMMA, GEMMA LOCO and the subpopulation meta-analysis. We assessed the ability of these methods to detect cis-eQTLs, as our approaches do not have power to detect trans-eQTLs in this simulation. Recall that the two simulation settings with cis effects assume that some large proportion of the variance is attributable to cis effects; thus, power differs for cis and trans effects in this simulation. We considered results at the transcript level, as we are not investigating fine mapping approaches that account for the effect of linkage disequilibrium. That is, if at least 1 significant cis-SNP is identified in a gene that has at least 1 truly nonzero cis effect, we count that as a true positive. That is the case even if the associated SNP is not identical to the truly nonzero SNP. Defining other events (false positive, false negative, true negative) follows from this logic. For example, a false positive occurs when we identify one or more associated cis-SNPs with our approach, but there is no truly associated cis-SNP. As previously, a SNP is identified as significant if its association test p-value is less than 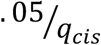.

Power and type I error (i.e., false positive rate) for the three analysis methods are assessed in the 43-strain setting. These results indicate that standard GEMMA and GEMMA LOCO are appropriate methods for the analysis of the HRDP data, while the subpopulation meta-analysis approach is substantially underpowered and thus suboptimal. First, we detail type I error and power results from the 43-strain simulation, then discuss the power of GEMMA LOCO applied to the subpopulation simulation since it was the method with the highest power in general. In eQTL studies, statistical power varies between transcript because of cis-heritability therefore, power is estimated separately for different ranges of cis-heritability.

The no cis effect simulation, in which cis and trans SNPs are simulated from the same distribution, is used to calculate the empirical type I error rate. In the 43-strain setting, each of the three methods has a type I error rate less than .015 (Supplementary Table 2). This is evidence that the type I error rate is well controlled by all of the methods; indeed, this indicates that the methods are somewhat conservative. This may be because we account for multiplicity in the cis region assuming that association tests for SNPs are independent, when in truth they are correlated due to linkage disequilibrium.

The one and multiple cis effect simulations in the 43-strain setting are used to assess the power of the analysis methods. The 43-strain HRDP has greater than 80% power to detect cis effects via the standard GEMMA and GEMMA LOCO approaches when the cis heritability exceeds 0.4, and reasonable power to detect cis effects when the cis heritability is between 0.1 and 0.4 (Figure 6). In the one cis-effect simulation, 73.4% of genes have cis-heritability larger than 0.1 and 43.5% of genes have cis-heritability larger than 0.4. In the multiple cis-effect simulation, 73.7% of genes have cis-heritability larger than 0.1 and 43.6% of genes have cis-heritability larger than 0.4. The standard GEMMA and GEMMA LOCO approaches have similar power, although GEMMA LOCO has slightly better power. The subpopulation meta-analysis approach has substantially less power to detect cis effects (Table 2). This may be because the quality control process for the subpopulation meta-analysis is more stringent, thus reducing the number of SNPs analyzed. This result indicates that the subpopulation meta-analysis approach is likely not optimal for analysis of the HRDP data.

**Figure 6.**
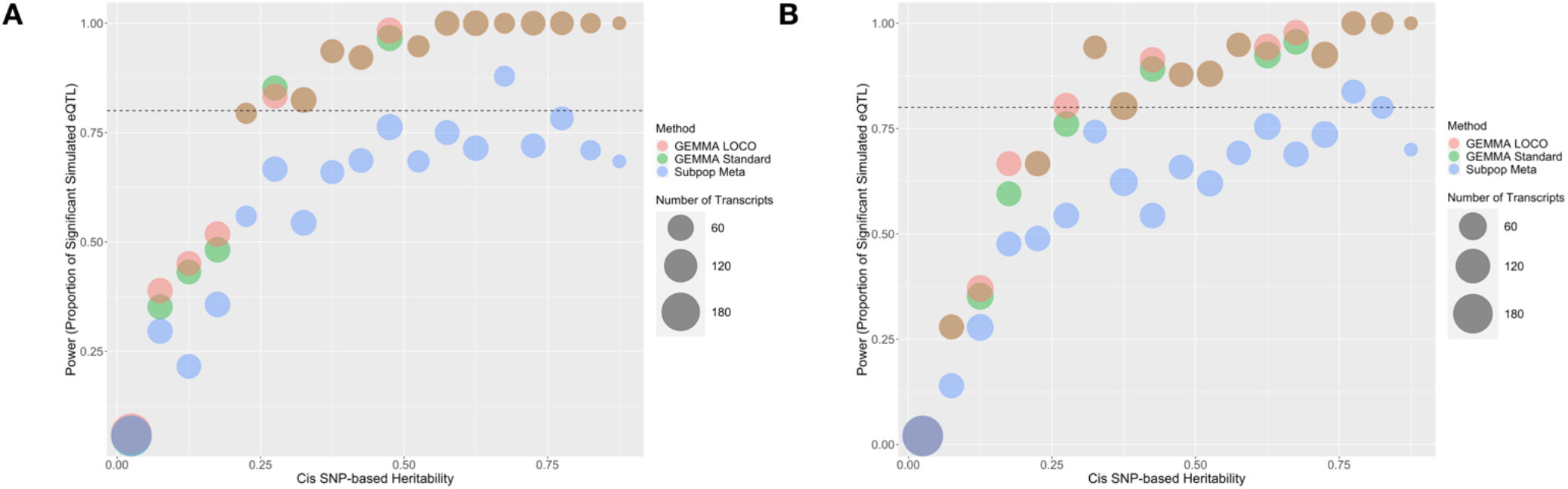
Predicted power in cis-eQTL studies based on cis-heritability of the transcript and QTL mapping method. The number of genes in each heritability strata is represented by the size of the bubble. Standard GEMMA is in green, GEMMA LOCO is in red, and subpopulation meta is in blue. Two simulation approaches were used that were based on either **A) a single cis-SNP effect** or **B) multiple cis-SNP effects**.

**Table 2.**
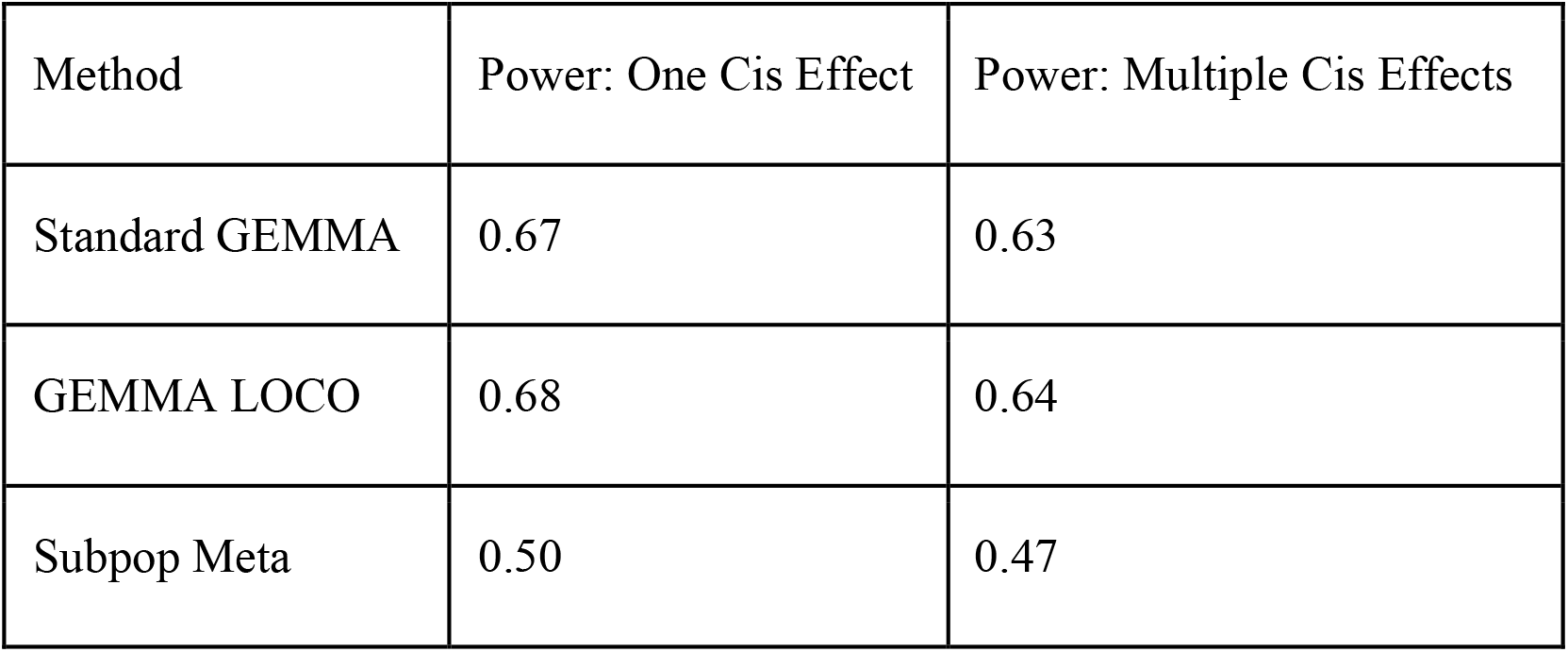
Power for standard GEMMA, GEMMA LOCO, and subpopulation meta-analysis applied to the two simulation settings with cis effects in the 43 strain simulations. Power is the proportion of genes with at least one nonzero cis-SNP effect that had at least one cis-SNP with a significant association identified by the corresponding model.

The 43-strain setting results indicate that standard GEMMA, GEMMA LOCO, and subpopulation meta-analysis are all appropriate for the analysis of the HRDP data, although the subpopulation meta-analysis is underpowered in comparison. Next, we applied GEMMA LOCO to four populations: the FXLE/LEXF recombinant inbred population, the HXB/BXH recombinant inbred population, the classic inbred population, and 92 strains of the HRDP. These results may inform study design decisions and how to select subpopulations from the HRDP for an eQTL study.

GEMMA LOCO has adequate type I error control when applied to all four populations, as shown in Supplementary Table 3. Figure 7 plots the empirical power of GEMMA LOCO applied to the four populations against the cis-heritability of the gene for the two simulation settings with cis-SNP effects. These results indicate that we have greater than 80% power to detect a cis-SNP effect using the 92 strains of the HRDP when the cis-heritability exceeds 0.25. GEMMA LOCO has similar power when applied to the classic inbred and HXB/BXH populations, although there is no cutoff point for cis-heritability after which we are guaranteed to exceed 80% power in both simulations. The power is notably worse for the FXLE/LEXF population despite its largest sample size among the three subpopulations (Table 3). As demonstrated by the kinship matrix in Figure 1, the FXLE/LEXF recombinant inbred panel is comprised of highly related strains, which reduces the effective sample size.

**Table 3.** Power for GEMMA LOCO applied to the two simulation settings with cis effects in the subpopulation simulations. Power is the proportion of genes with at least one nonzero cis-SNP effect that had at least one cis-SNP with a significant association identified via GEMMA LOCO analysis of the corresponding subpopulation.

| Subpopulation | Power: One Cis Effect | Power: Multiple Cis Effects |
| --- | --- | --- |
| FXLE/LEXF (33 strains) | 0.25 | 0.30 |
| HXB/BXH (30 strains) | 0.53 | 0.52 |
| Classic Inbred (29 strains) | 0.51 | 0.52 |
| HRDP (92 strains) | 0.74 | 0.74 |

**Figure 7.**
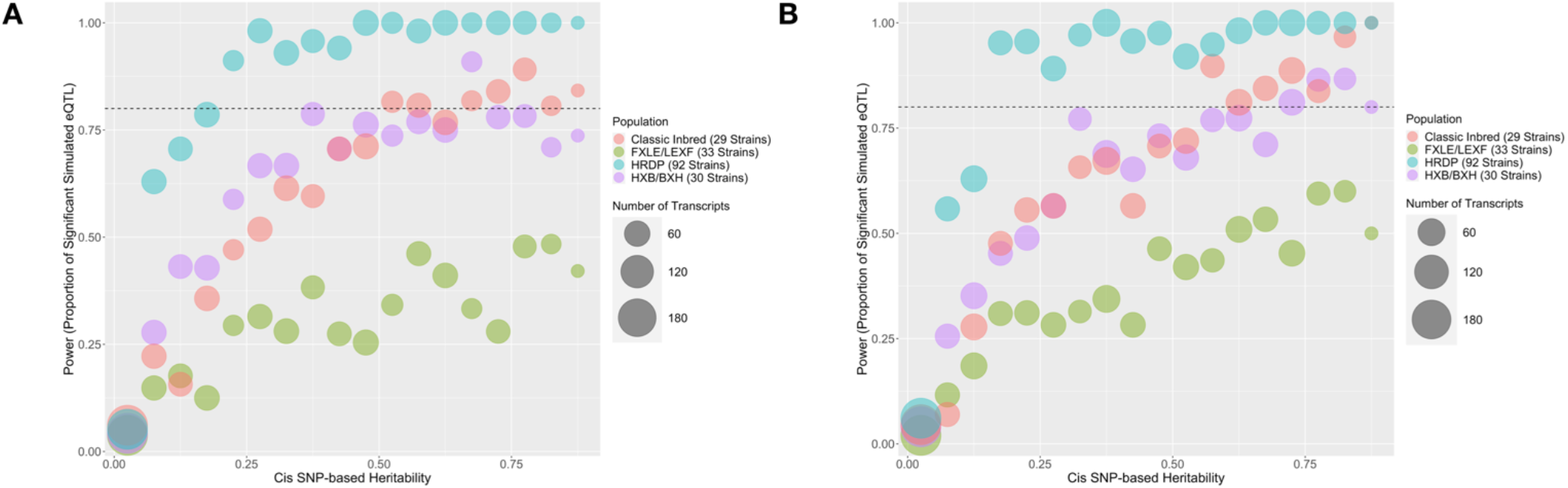
Predicted power in cis-eQTL studies based on cis-heritability of the transcript and rat population used for QTL mapping. The number of genes in each heritability strata is represented by the size of the bubble. The HXB/BXH RI population is in purple, the FXLE/LEXF RI population is in green, the classic inbred population is in red, and the full HRDP is in blue. Two simulation approaches were used that were based on either **A) a single cis-SNP effect** or **B) multiple cis-SNP effects**.

## 4 Discussion

This paper discusses several feasible methods for eQTL analyses in the Hybrid Rat Diversity Panel. In particular, we rigorously investigate the performance of four approaches to QTL analysis: linear regression, linear mixed model, linear mixed model with leave one chromosome out kinship matrix calculation, and linear mixed model by subpopulation followed by meta-analysis. We apply these methods to brain RNA expression levels in 99,376 transcripts in the HRDP and characterize the performance of these methods via genomic inflation and the number of associated cis-SNPs. We further explore the type I error and power of these methods via simulation. Finally, we characterize the power of QTL analyses in subpopulations of the HRDP data, and empirically relate the power to the phenotypic heritability. We conduct the power analyses by subpopulation and for the entire set of 92 strains to facilitate study design decisions regarding the HRDP data.

From this analysis, we conclude that the standard GEMMA and GEMMA LOCO approaches both control type I error sufficiently and maximize power. Therefore, both are feasible choices for eQTL analyses of the HRDP data. GEMMA LOCO identifies the most significant cis-SNPs in the real data application and has the best power in simulation while maintaining the nominal level of type I error control in simulation. The genomic inflation incurred by the GEMMA LOCO analysis of the real data is some evidence that it may have weaker type I error control than the other approaches. For applications where controlling type I error is of great emphasis, we believe that it is also appropriate to use the standard GEMMA approach, as this approach is only marginally less powerful than GEMMA LOCO.

Figures 6 and 7 display the power of the analysis methods applied to different subpopulations within the HRDP. These plots may be useful for future study design, in particular for determining sample size and deciding which subpopulation of the HRDP data to use. For researchers interested in using only one of the three subpopulations within the HRDP, it appears that the FXLE/LEXF recombinant inbred subpopulation will yield the least power, while the HXB/BXH recombinant inbred panel and the set of classic inbred strains have similar power. Applying GEMMA LOCO to a set of 92 HRDP strains is quite powerful; per our simulation, we have >80% power to detect an association when the cis-heritability of a candidate gene exceeds .15.

These results establish a baseline of modeling recommendations and study design considerations for eQTL analyses of the HRDP. Given the central role transcript variation plays in the regulation of complex disease and the substantial effect of genetic variation on transcript variation, characterizing the genetic control of transcript variation is of great interest. Transcriptome analysis in human subjects with complex disease is often confounded by substantial environmental variation, making model organism analysis an attractive alternative for further parsing genetic control of the transcriptome. We hope this study will facilitate rigorous statistical analyses of well-powered studies involving the HRDP, thus furthering systems genetics applications towards the understanding of complex disease.

## 5 Author Contributions

HRDP database study design: BT, PH, and LS. Data processing: JP, LS, SM, and LV. Statistical analysis: JP and LV. Writing: JP, LV, and LS with input from all authors.

## 8 Acknowledgments

The authors thank Jennifer Mahaffey for technical support on the project.

## 9 Data Availability Statement

The brain transcriptome data analyzed in this study is available through the NCBI GEO browser, accession number GSE199984. The processed HRDP brain RNA expression data and the genotype data is available for download through PhenoGen (https://phenogen.org/publication/pattee_2022).

