## Supplementary material for "Power and precision: Evaluation and recommendations of quantitative trait analysis methods for RNA expression levels in the Hybrid Rat Diversity Panel"

April 2022

#### S1 Simulation Study Design

This supplementary section describes the design of the simulation study and some preliminary results involving the simulated data. We extensively detail the simulation method and present some results indicating that the simulated data coheres well to the real data.

A general overview of the simulation setup is as follows. We used real genetic marker data from 92 HRDP subjects genotyped on 18,342 SNPs in the simulation. After simulating SNP effects in the manner described throughout this supplementary section, we simulated phenotypes by matrix multiplication between the real genotype data and the simulated SNP effects. While data are simulated using the entire set of 18,342 SNPs, only SNPs that pass the quality control described in the manuscript (i.e., minor allele frequency $> .1$, missinginess $< 5\%$) are included as fixed effects in the inferential model. Broadly, there are two analyses we are interested in: the set of 43 subjects for which we have real data ('43-subject simulations') and the full set of 92 subjects and the three subpopulations ('subpopulation simulations'). In the 43-subject simulation, we apply the GEMMA LOCO, standard GEMMA, and subpopulation meta-analysis methods. In the subpopulation simulations, we apply only GEMMA LOCO. The intention of the 43-subject simulations is to establish the power and type I error of the three competing methods. The intention of the subpopulation simulations is to compare the power of GEMMA LOCO to detect SNP effects in the full HRDP panel to each of the subpopulations.

The simulation study involved three unique simulation settings, as follows:

1. 'No cis effect' or 'unimodal' simulation, where there is no difference in effect size between cis SNPs and trans SNPs.
2. 'One cis effect' or 'bimodal one cis' where cis SNPs contribute to a substantially larger proportion of the heritability than trans SNPs, and exactly 1 cis-SNP has nonzero effect size.
3. 'Multiple cis effects' or 'bimodal multiple cis', where cis-SNPs contribute to a substantailly larger proportion of the heritability than trans SNPs, and the number of cis-SNPs with nonzero effect size is simulated from a binomial distribution, with mean number of nonzero cis-SNPs equal to two.

These three simulation protocols are used to generate simulated phenotypes for the full set of 92 HRDP subjects. The subsequent analyses, i.e. the 43-subject simulations and the subpopulation simulations, are conducted by appropriately subsampling from the full genotype and simulated phenotype data. For the 43 subject simulations, we apply the standard GEMMA, GEMMA LOCO, and subpopulation meta-analysis methods to each of the three simulation protocols, which is a total of 9 analyses. For the subpopulation simulations, we apply the GEMMA LOCO model to each simulation setting in each of the following four populations: all 92 subjects, only recominant HXB/BXH, only recominant FXLE/LEXF, only classic inbred. This is a total of 12 analyses. Table S1 describes the full set of 21 analyses conducted.

| | No cis effect | One cis effect | Multiple Cis Effect |
|---|---|---|---|
| 43-Subject | GS, GL, SM | GS, GL, SM | GS, GL, SM |
| Full HRDP (92 Strains) | GL | GL | GL |
| HXB/BXH (30 Strains) | GL | GL | GL |
| FXLE/LEXF (33 Strains) | GL | GL | GL |
| Classic Inbred Only (29 Strains) | GL | GL | GL |

**Table S1:** Analyses performed for each population and each simulation setting. **GS** signifies 'GEMMA Standard', **GL** signifies 'GEMMA LOCO', **SM** signifies 'Supopluation meta-analysis'

Each simulated gene expression was simulated to have heritability equal to the (GEMMA estimated) heritability of the gene from the real data. The correspondence is, of course, not exact due to stochasticity. For the no cis effect simulation, we used only genes estimated to be unimodal in the real data. For the two bimodal simulations, we used only genes estimated to be bi- or tri-modal in the real data. In the following three subsections, we present some results to confirm that our setup is working as intended. The following three subsections describe the three simulation settings in detail.

#### S1.1 No cis effect simulation

For the unimodal simulation, we simulate SNP effects from a point-normal distribution as follows, for all $j \in [1, \ldots, 18, 342]$ SNPs:

$$\beta_j \sim_{iid} \begin{cases} N(0, \frac{h^2}{p \times M}), \text{ with probability p} \\ 0, \text{ with probability 1-p} \end{cases}$$

We fix $p = 1/10$, which entails that 10% of SNPs have a heritable nonzero contribution to the phenotype. $M$ is set equal to the number of SNPs (18,342). $h^2$ is the chip-based hertiability from the gene estimated via standard GEMMA on the real data. We simulated phenotypes for a random sample of 1000 genes drawn from the 81,992 genes that were estimated to be unimodal in the real data. After simulating SNP effects from the above point-normal model, we have the heritable portion of the phenotype, which we can denote as $\mathbf{X}\boldsymbol{\beta}$. Note here $\mathbf{X}$ is the genotype matrix of SNP data. We then simulate error variance according to the following distribution: $\boldsymbol{\epsilon} \sim N(0, \frac{(1-h^2) \times var(\mathbf{X}\boldsymbol{\beta})}{h^2}\mathbf{I})$ and define our gene expression phenotype $\mathbf{y} = \mathbf{X}\boldsymbol{\beta} + \boldsymbol{\epsilon}$. We uniquely simulate phenotypes for 1000 genes in this manner.

We are interested in whether the simulated data coheres well to the real data, which is investigated as follows. Firstly, we see how well simulating in this manner generates a distribution of heritability that matches the true per-gene heritability. Figure S1 shows that the simulated heritability (calculated here as $cor(\mathbf{X}\boldsymbol{\beta}, \mathbf{y})^2$) is similar to the true heritability. Secondly, we investigate whether simulating gene expression in this fashion produces gene expression phenotypes that are truly unimodal. We used the R package mclust to estimate modality for the 43-subject simulation in the manner as for the real data, i.e. fitting uni, bi, and trimodal models and selecting the optimal model via BIC. Figure S2 displays density plots and histograms for three randomly chosen genes, and table S2 displays a count of the number of genes with each modal assignment. These results indicate that this simulation setting does a reasonable job of inducing unimodality in the simulated phenotype.

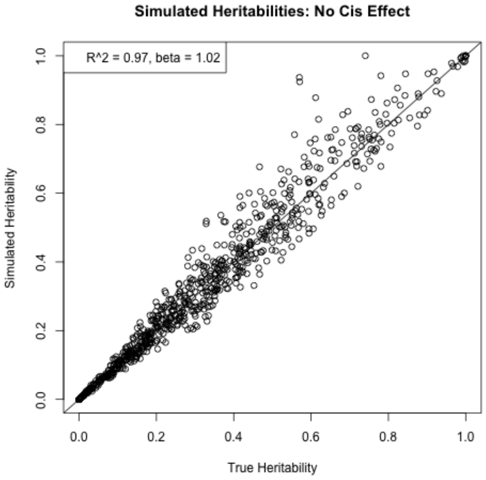


**Figure S1:** Simulated chip-based heritability versus true chip-based heritability for 1000 simulated unimodal genes.

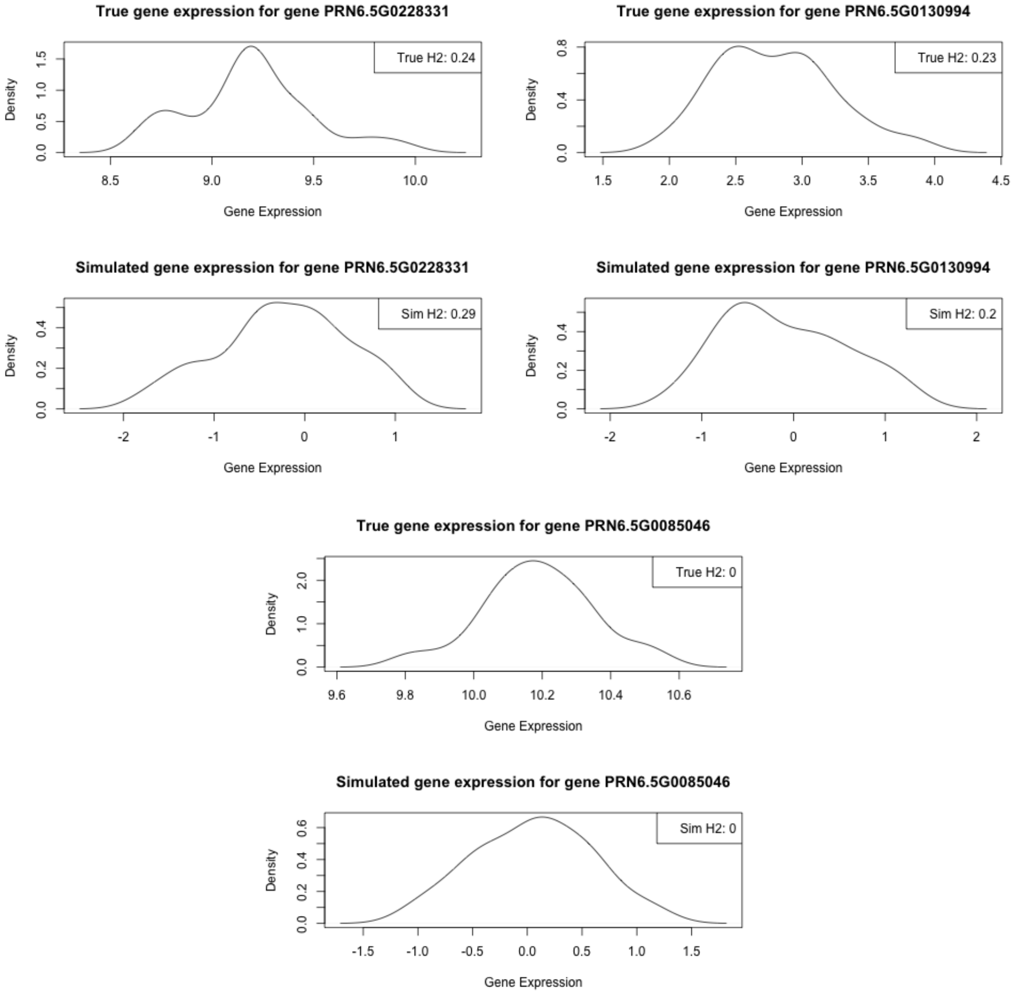


**Figure S2:** Distribution of simulated versus true gene expression for three randomly chosen unimodal genes (out of 1000).

| Cluster Assignments | Unimodal | Bimodal | Trimodal |
|---|---|---|---|
| Count of Genes | 954 | 42 | 4 |

**Table S2:** Count of genes per number of modes as estimated by 'mclust', for unimodal simulation.

#### S1.2 One cis effect simulation

For the one cis effect simulation, we simulate phenotypes for a randomly chosen set of 1,000 genes out of the 17,384 genes that are estimated to be bimodal or trimodal in the real data. To induce bimodality in our simulated phenotypes, we simulate SNP effects for the cis and trans regions from different point-normal distributions. We fix the proportion of the total heritability attributable the cis region at $T$ (denoting 'heritability Tradeoff'). For each of the 1000 genes that we simulate, we draw $T$ from a $Unif(.7, .9)$ distribution. Consider that we have SNPs indexed by $i \in [1, \ldots, q_{cis}]$, where $q_{cis}$ is the number of cis-SNPs. Let $s$ be some randomly chosen index such that $1 \leq s \leq q_{cis}$. We then define the point-normal distribution for the cis-SNPs as follows:

$$\beta_j \sim_{iid} \begin{cases} N(0, \frac{h^2_{cis}}{p \times q_{cis}}), \text{for SNP} j = s \\ 0, \text{for all SNPs } j \text{ such that } j \neq s \end{cases}$$

In this way, we ensure that exactly one (randomly chosen) cis-SNP has nonzero effect size. We define $p = 1/q_{cis}$. $h^2_{cis}$ is defined as the true heritability of the gene (estimated from real data) multiplied by $T$: that is, $h^2_{cis} = T \times h^2$.

Defining the distribution for the trans-SNPs is more difficult. Ultimately, we want that $h^2_{trans} + h^2_{cis} = h^2$. In order for this expression to hold, we need to ensure that the heritability contributed from the cis-SNPs and the trans-SNPs is orthogonal. To ensure this, we only simulate nonzero effects for trans-SNPs that are on different chromosomes than the gene in question. That is, all trans SNPs on the same chromosome as the gene have effect set equal to zero. We also want that $\frac{h^2_{cis}}{h^2} = T$. If we partition $\boldsymbol{\beta}$ into cis and trans components $\boldsymbol{\beta}_{trans}$ and $\boldsymbol{\beta}_{cis}$ and do likewise with the genotype matrix $\mathbf{X}_{trans}$ and $\mathbf{X}_{cis}$, we can define the cis heritability as follows:

$$h^2_{cis} = \frac{var(\mathbf{X}_{cis}\boldsymbol{\beta}_{cis})}{var(\mathbf{X}_{cis}\boldsymbol{\beta}_{cis}) + var(\mathbf{X}_{trans}\boldsymbol{\beta}_{trans}) + var(\boldsymbol{\epsilon})}$$

Here, we see the difficulty in defining a distribution for the trans effects: namely, that the variance of $\mathbf{X}_{trans}\boldsymbol{\beta}_{trans}$ depends on the distribution of the specific genotype matrix $\mathbf{X}_{trans}$ for that gene. We approach this as follows. We initialize the trans effects according to the point normal distribution as follows:

$$\beta_j \sim_{iid} \begin{cases} N(0, \frac{h^2_{trans}}{p \times q_{trans}}), \text{ with probability p} \\ 0, \text{ with probability 1-p} \end{cases}$$

With $p = 1/10$ and $q_{trans}$, defined as the count of all SNPs not on the same chromosome as the gene in question. Given this variance, we simulate SNP effects, and calculate $var(\mathbf{X}_{trans}\boldsymbol{\beta}_{trans})$. If the following expression is satisfied:

$$var(\mathbf{X}_{trans}\boldsymbol{\beta}_{trans}) < \frac{(1 - T) \times var(\mathbf{X}_{cis}\boldsymbol{\beta}_{cis})}{T}$$

We use the simulated $\boldsymbol{\beta}_{trans}$. Otherwise, we divide the variance of the effect size distribution $\frac{h^2_{trans}}{Mp}$ by 2, and simulate trans effects again until the condition is satisfied.

Once this is done, we can simulate normal errors such that the following expression holds:

$$h^2 = \frac{var(\mathbf{X}_{cis}\boldsymbol{\beta}_{cis}) + var(\mathbf{X}_{trans}\boldsymbol{\beta}_{trans})}{var(\mathbf{X}_{cis}\boldsymbol{\beta}_{cis}) + var(\mathbf{X}_{trans}\boldsymbol{\beta}_{trans}) + var(\boldsymbol{\epsilon})}$$

Figure S3 demonstrates that simulating effect sizes in the specified manner generates genes with roughly the desired heritability. Figure S4 displays the distribution of the simulated gene expression for three randomly selected bimodal genes versus the true distribution of the gene expression. Table S3 displays the estimated modality of the simulated phenotypes for the 43-subject simulation. We see that this simulation setting induces some amount of bimodality, but a majority of genes are estimated to be unimodal.

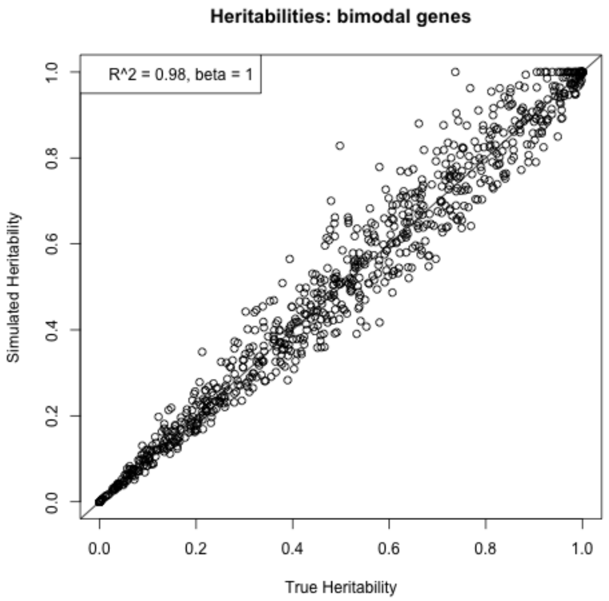


**Figure S3:** Simulated chip-based heritability versus true chip-based heritability for 1000 simulated bimodal genes (simulation 2).

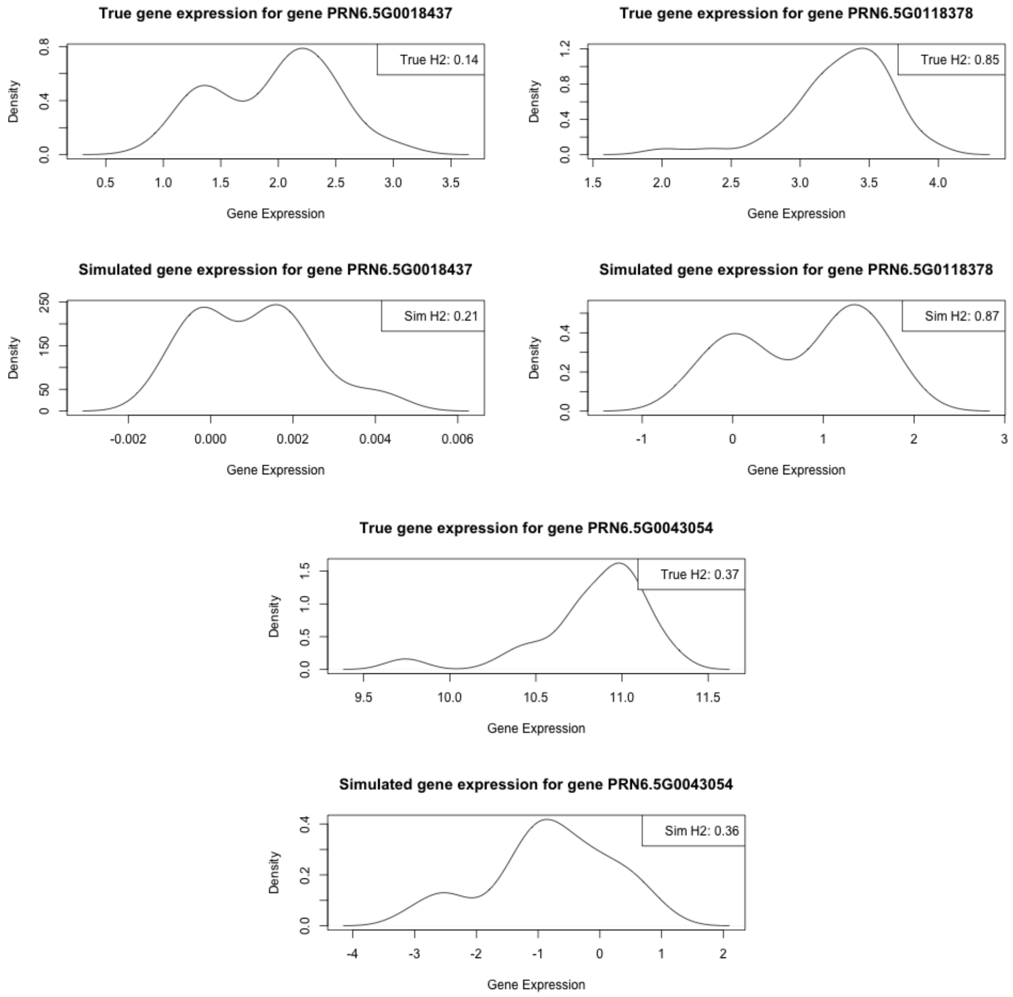


**Figure S4:** Distribution of simulated versus true gene expression for three randomly chosen bimodal genes (simulation 2).

| Cluster Assignments | Unimodal | Bimodal | Trimodal |
|---|---|---|---|
| Count of Genes | 760 | 222 | 18 |

**Table S3:** Count of genes per number of modes as estimated by 'mclust', for bimodal simulation (simulation 2).

#### S1.3 Multiple cis effect simulation

The setup of the multiple cis effect simulation is very similar to the setup of the single cis effect simulation, so many details will not be recapitulated here. The distinction is that the cis-SNPs are simulated from a different point-normal distribution, which is as follows:

$$\beta_j \sim_{iid} \begin{cases} N(0, \frac{h^2_{cis}}{p \times M_{cis}}), \text{ with probability p} \\ 0, \text{ with probability 1-p} \end{cases}$$

We fix $p$ so that the expected number of cis-SNPs with nonzero effects is 2. That is, given $M_{cis}$ which denotes the number of cis-SNPs in the gene, we define $p = 2/M_{cis}$. The rest of the quantities in the expression are the same as described in section **S1.2**.

Figure S5 shows that simulating effect sizes according to the above process does produce the desired heritabilities with no systemic bias. Figure S6 plots the distribution of the true and simulated phenotype for three randomly selected genes. Table S4 displays the estimated modality of the simulated phenotypes for the 43-subject simulation. Again, the simulation induces a reasonable amount of bimodality, but a majority of the genes are estimated to be unimodal.

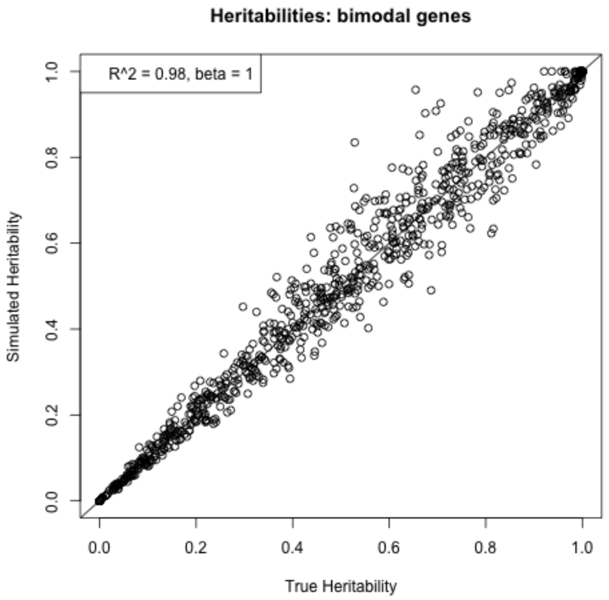


**Figure S5:** Simulated chip-based heritability versus true chip-based heritability for 1000 simulated bimodal genes (simulation 3).

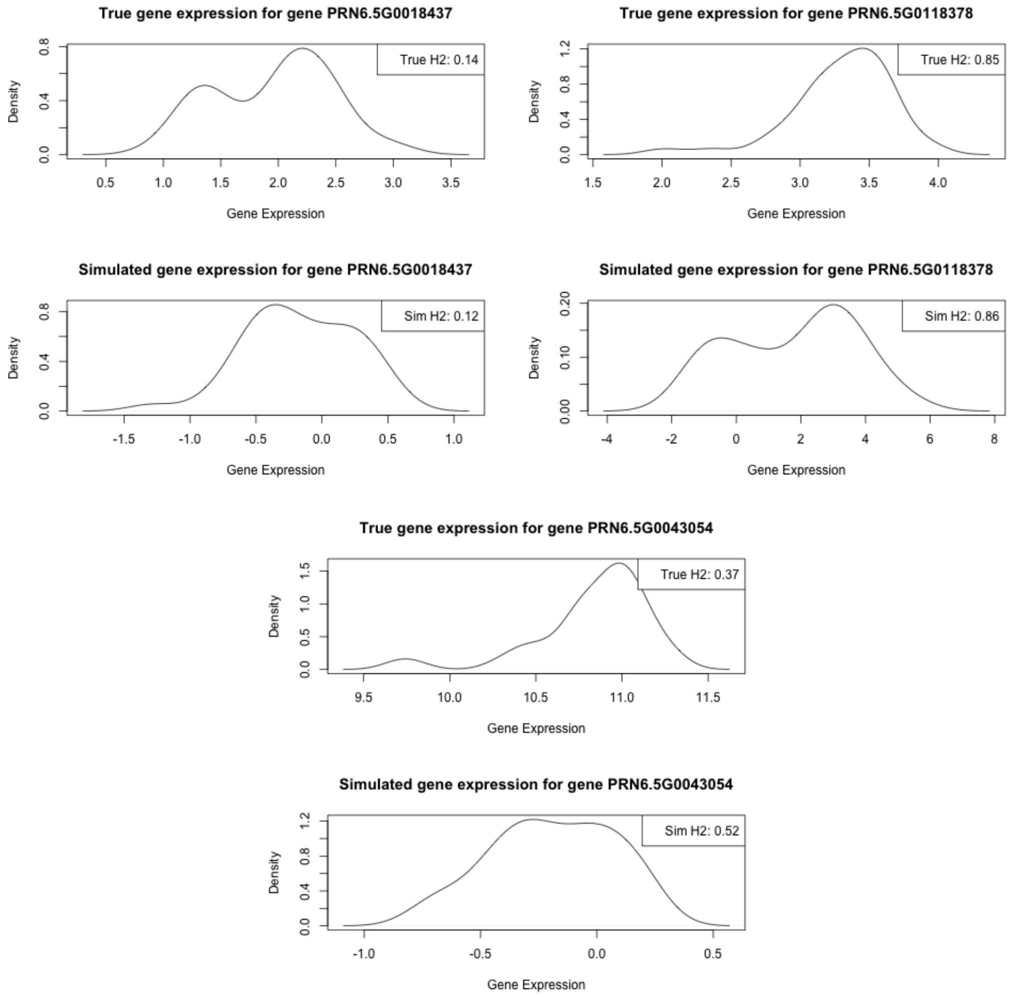


**Figure S6:** Distribution of simulated versus true gene expression for three randomly chosen bimodal genes (simulation 3).

| Cluster Assignments | Unimodal | Bimodal | Trimodal |
|---|---|---|---|
| Count of Genes | 703 | 290 | 7 |

**Table S4:** Count of genes per number of modes as estimated by 'mclust', for bimodal simulation (simulation 3).

#### S2 Supplementary Figures

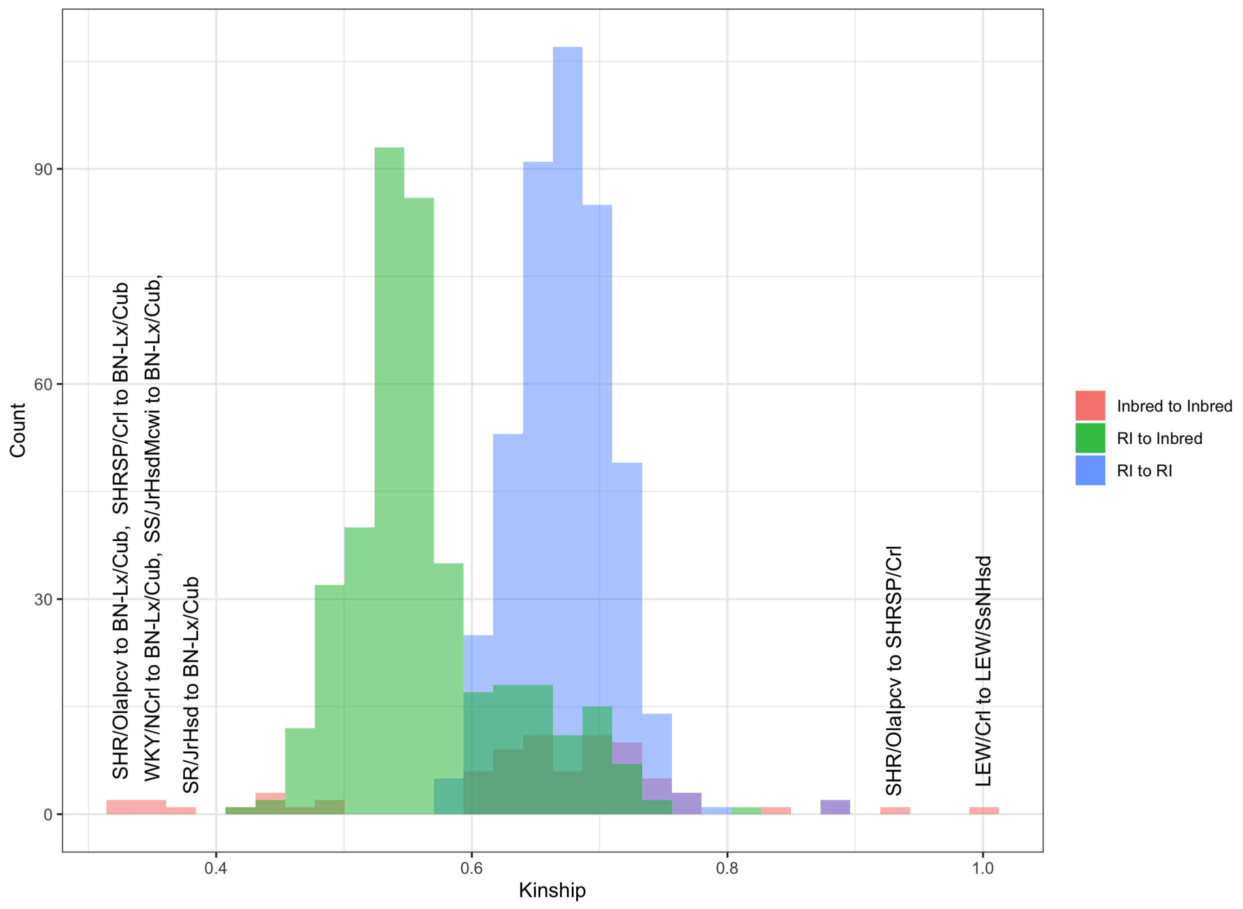


**Supplementary Figure 1:** Kinship values for each pairwise relationship between strains. Red denotes inbred to inbred kinship values, blue denotes RI to inbred kinship values and green denotes RI to RI kinship values. Extreme values are labeled with their corresponding pairs.

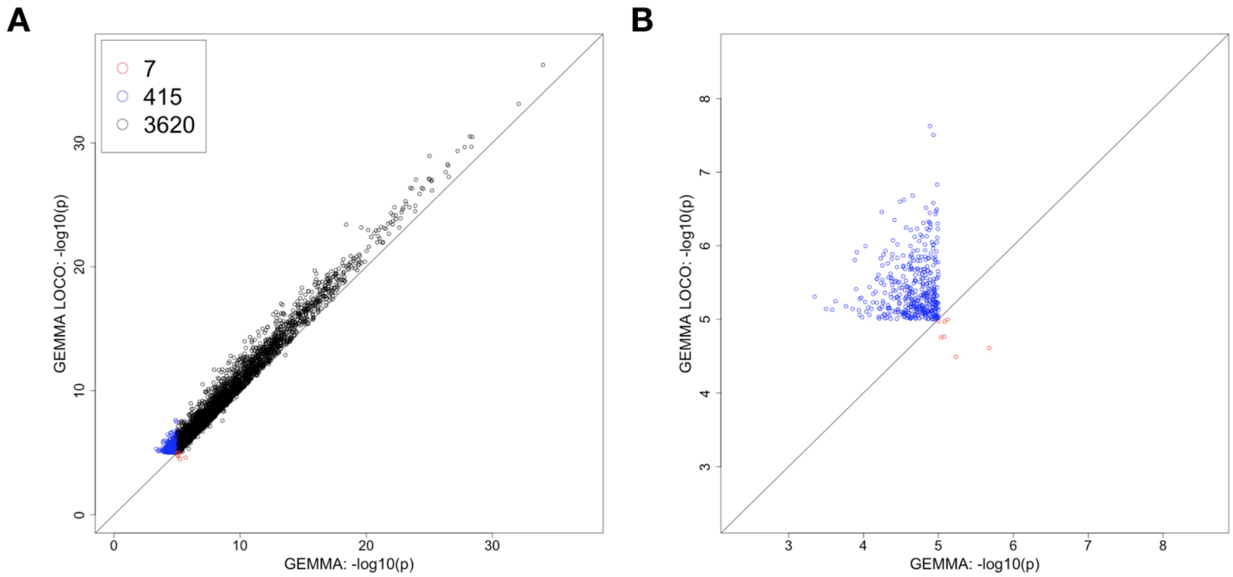


**Supplementary Figure 2:** Comparison of the negative base 10 logarithm of the p value of the most significant cis-SNP for the standard GEMMA (x axis) and GEMMA LOCO (y axis) models. Each point on the plot represents a gene with at least one cis-SNP identified as significant by at least one of the two methods. Panel A) shows associations that are significant in only GEMMA LOCO (blue), only GEMMA standard (red), and both (black). Panel B) shows only associations that are significant in exactly one analysis.

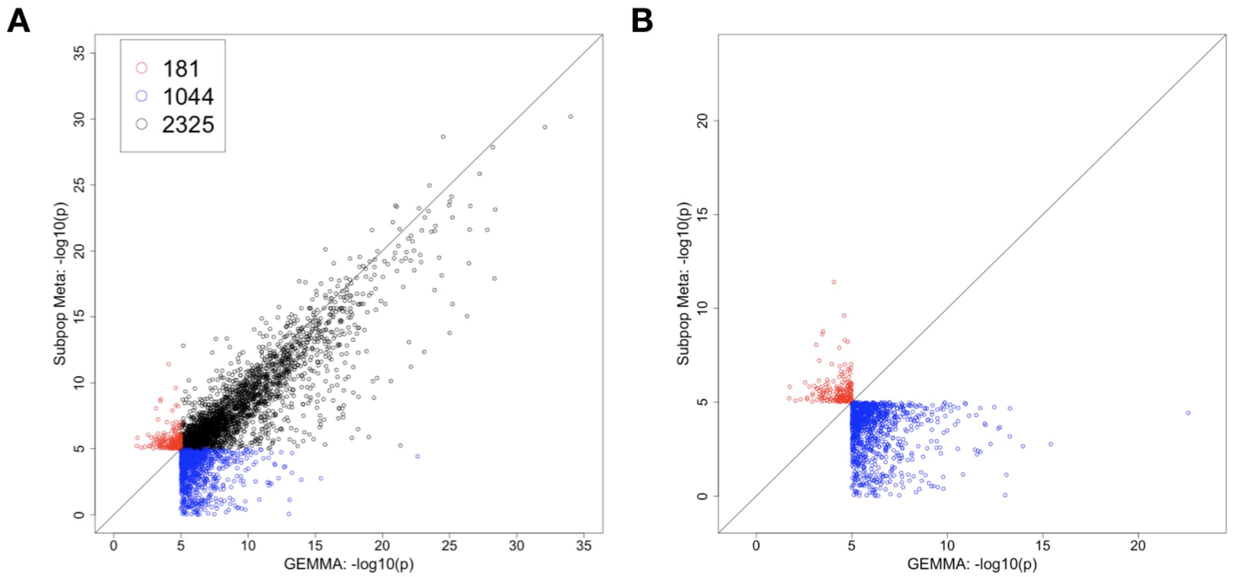


**Supplementary Figure 3:** Comparison of the negative base 10 logarithm of the p value of the most significant cis-SNP for the standard GEMMA (x axis) and subpopulation meta (y axis) models. Each point on the plot represents a gene with at least one cis-SNP identified as significant by at least one of the two methods. Panel A) shows associations that are significant in only GEMMA (blue), only in subpopulation meta (red), and both (black). Panel B) shows only associations that are significant in exactly one analysis.

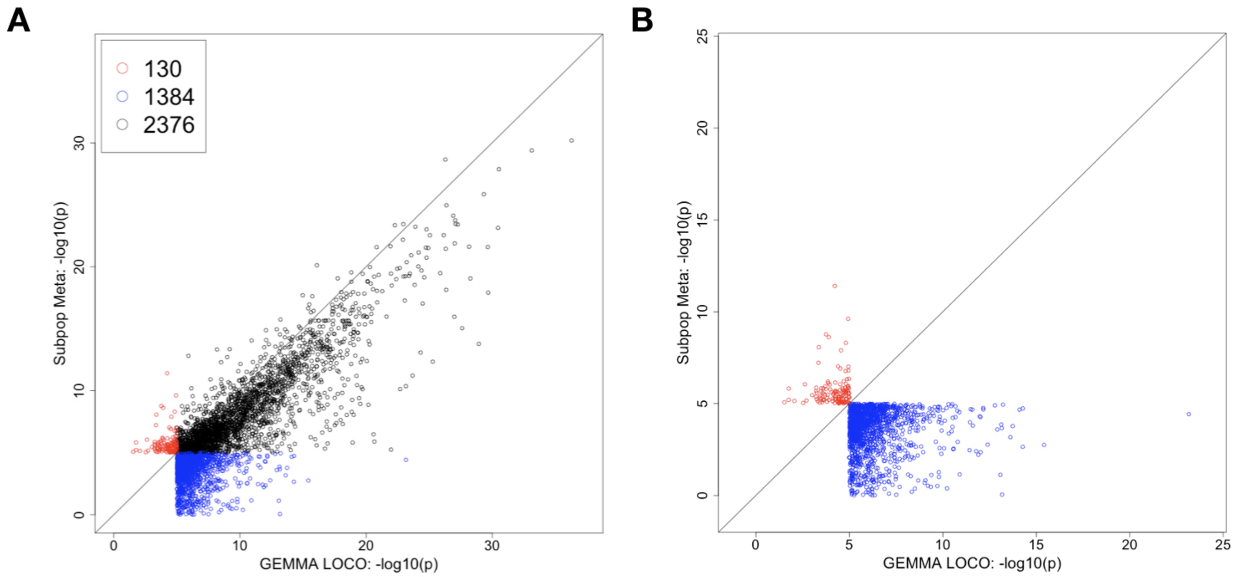


**Supplementary Figure 4:** Comparison of the negative base 10 logarithm of the p value of the most significant cis-SNP for the GEMMA LOCO (x axis) and subpopulation meta (y axis) models. Each point on the plot represents a gene with at least one cis-SNP identified as significant by at least one of the two methods. Panel A) shows associations that are significant in only GEMMA LOCO (blue), only in subpopulation meta (red), and both (black). Panel B) shows only associations that are significant in exactly one analysis.

### S3 Supplementary Tables

| | Minimum | 1st Quartile | Median | Mean | 3rd Quartile | Maximum |
|---|---|---|---|---|---|---|
| GEMMA | 0.34 | 1.00 | 1.13 | 1.12 | 1.25 | 2.45 |
| GEMMA LOCO | 0.34 | 1.07 | 1.22 | 1.19 | 1.33 | 2.27 |
| Subpopulation Meta | 0.45 | 0.98 | 1.09 | 1.10 | 1.20 | 2.66 |
| Linear Regression | 1.94 | 3.01 | 3.78 | 4.48 | 5.34 | 15.08 |

**Supplementary Table 1:** The distribution of the genomic inflation factor across 99,736 eQTL transcripts for each of the four methods.

| Method | Type I Error |
|---|---|
| GEMMA | 0.008 |
| GEMMA LOCO | 0.010 |
| Subpopulation Meta | 0.015 |

**Supplementary Table 2:** Type I error for the three methods applied to the unimodal simulation for the 43-strain setting.

| Subpopulation | Type I Error |
|---|---|
| FXLE/LEXF | 0.014 |
| HXB/BXH | 0.025 |
| Classic Inbred | 0.037 |
| All 92 | 0.041 |

**Supplementary Table 3:** Type I error for GEMMA LOCO analyses of the four subpopulations in unimodal simulation.
